# PET Imaging of the Serotonin 1A Receptor in Major Depressive Disorder: Hierarchical Multivariate Analysis of [^11^C]WAY100635 Overcomes Outcome Measure Discrepancies

**DOI:** 10.1101/2024.03.12.584569

**Authors:** Granville J. Matheson, Francesca Zanderigo, Jeffrey M. Miller, Elizabeth A. Bartlett, J. John Mann, R. Todd Ogden

## Abstract

The serotonin 1A receptor has been linked to both the pathophysiology of major depressive disorder (MDD) and the antidepressant action of serotonin reuptake inhibitors. Most PET studies of the serotonin 1A receptor in MDD used the receptor antagonist radioligand, [carbonyl-^11^C]WAY100635; however the interpretation of the combined results has been contentious owing to reports of higher or lower binding in MDD with different outcome measures. The reasons for these divergent results originate from several sources, including properties of the radiotracer itself, which complicate its quantification and interpretation; as well as from previously reported differences between MDD and healthy volunteers in both reference tissue binding and plasma free fraction, which are typically assumed not to differ. Recently, we have developed two novel hierarchical multivariate methods which we validated for the quantification and analysis of [^11^C]WAY100635, which show better accuracy and inferential efficiency compared to standard analysis approaches. Importantly, these new methods should theoretically be more resilient to many of the factors thought to have caused the discrepancies observed in previous studies. We sought to apply these methods in the largest [^11^C]WAY100635 sample to date, consisting of 160 individuals, including 103 MDD patients, of whom 50 were not-recently-medicated and 53 were antidepressant-exposed, as well as 57 healthy volunteers. While the outcome measure discrepancies were substantial using conventional univariate analysis, our multivariate analysis techniques instead yielded highly consistent results across PET outcome measures and across pharmacokinetic models, with all approaches showing higher serotonin 1A autoreceptor binding potential in the raphe nuclei of not-recently-medicated MDD patients relative to both healthy volunteers and antidepressant-exposed MDD patients. Moreover, with the additional precision of estimates afforded by this approach, we can show that while binding is also higher in projection areas in this group, these group differences are approximately half of those in the raphe nuclei, which are statistically distinguishable from one another. These results are consistent with the biological role of the serotonin 1A autoreceptor in the raphe nuclei in regulating serotonin neuron firing and release, and with preclinical and clinical evidence of deficient serotonin activity in MDD due to over expression of autoreceptors resulting from genetic and/or epigenetic effects. These results are also consistent with downregulation of autoreceptors as a mechanism of action of selective serotonin reuptake inhibitors. In summary, the results using multivariate analysis approaches therefore demonstrate both face and convergent validity, and may serve to provide a resolution and consensus interpretation for the disparate results of previous studies examining the serotonin 1A receptor in MDD.

## 1 INTRODUCTION

The serotonin (5-HT) 1A receptor (5-HT_1A_R) serves as a postsynaptic heteroreceptor in most of the brain, and as a presynaptic autoreceptor in the raphe nuclei where serotonin neuron cell bodies are located (Barnes and Sharp, 1999; Hoyer et al., 2002). Owing to its autoreceptor function, it is able to centrally regulate firing rates of serotonin neuron and the release levels of 5-HT throughout the brain. The 5-HT_1A_R has been implicated in the pathophysiology of several psychiatric and neurological disorders (Arango et al., 2001; Tiihonen et al., 2004; Savic et al., 2004; Kepe et al., 2006; Akimova et al., 2009), and of major depressive disorder (MDD) in particular (Boldrini et al., 2008; Kaufman et al., 2016). The autoreceptor is implicated in both the pathophysiology of MDD, as well as the antidepressant action of many antidepressant medications (Blier and De Montigny, 1983; Richardson-Jones et al., 2010; Gray et al., 2013). For these reasons, the 5-HT_1A_R has been and remains of substantial research interest and importance for human molecular imaging studies of MDD patients. In humans, *in vivo* quantification of 5-HT_1A_R is performed using PET, with most studies using [carbonyl-^11^C]WAY100635, hereon referred to as [^11^C]WAY100635, due to its high affinity, target selectivity and favorable kinetics (Farde et al., 1997; Osman et al., 1998; Kumar and Mann, 2007). Over the last two decades, numerous PET studies have examined [^11^C]WAY100635 binding comparing MDD patients and healthy volunteers (HV) —however results have been mixed, making their interpretation contentious (Parsey et al., 2010; Shrestha et al., 2012). The major source of this disagreement regarding the direction of the findings concerns the quantification of [^11^C]WAY100635 binding and the use of different outcomes measures across different studies (for more details, see Supplementary Materials S1).

The study of the 5-HT_1A_R using PET in humans has been particularly challenging due both to the properties of the [^11^C]WAY100635 radioligand and the neurobiology of the 5-HT_1A_R. The majority of studies have used reference tissue (i.e. indirect) approaches to estimate *BP*_ND_ using the cerebellum as a reference region, without collecting arterial plasma. However, the interpretation of these studies is complicated by the fact that the extremely low non-displaceable binding of [^11^C]WAY100635 renders the cerebellum disproportionately sensitive to the small degree of contamination from radiometabolites (Osman et al., 1998; Shrestha et al., 2012). Because the relationship of *BP*_ND_ with non-displaceable binding is proportional rather than subtractive (as it is for *BP*_P_ or *BP*_F_), indirect estimation of *BP*_ND_ is particularly prone to bias in cerebellar binding estimates, and this contamination is thought to produce inaccurate estimates. Moreover, studies have shown not only that there is a non-negligible degree of specific binding of [^11^C]WAY100635 in the cerebellar grey matter which differs between individuals (Parsey et al., 2005; Hirvonen et al., 2007; Parsey et al., 2010), but also that cerebellar binding is higher in MDD patients than healthy volunteers (HV) (Parsey et al., 2010). Together with the low non-displaceable binding of [^11^C]WAY100635 and the high sensitivity of *BP*_ND_ to small biases in cerebellar binding, group differences in cerebellar grey matter binding induce artificially lower estimates of *BP*_ND_ in MDD patients when it is used as a reference region (Parsey et al., 2010). Cerebellar white matter has therefore been recommended as the preferred reference region (Parsey et al., 2005; Hirvonen et al., 2007); however, this region is even more affected by radiometabolites due to its lower binding. On the basis of these issues, caution has been urged generally for the interpretation of *BP*_ND_ for the analysis of [^11^C]WAY100635 data (Parsey et al., 2005; Hirvonen et al., 2007; Parsey et al., 2010; Shrestha et al., 2012).

Indirect quantification of *BP*_P_ and *BP*_F_ using arterial plasma allows researchers to minimise the influence of these biases owing to the subtractive, as opposed to proportional, relationship of these outcomes with cerebellar binding. In the subset of studies in which arterial plasma samples were collected, results using *BP*_P_ have been mixed (Meltzer et al., 2004; Parsey et al., 2006; Hirvonen et al., 2008), while results using *BP*_F_ have been more consistent. Parsey et al. (2006) observed globally higher [^11^C]WAY100635 *BP*_F_ in unmedicated MDD patients compared to HV, which was later replicated in two subsequent studies (Parsey et al., 2010; Miller et al., 2013). However, interpretation of these studies is not straightforward either: although differences were observed using *BP*_F_, *f*_P_ differed significantly between MDD and control groups (Parsey et al., 2010). Reasons for group differences in *f*_P_ are not well understood, and it cannot be known whether these differences could have been caused by some experimental confound such as experimental drift, or whether they were biological. Unfortunately however, *BP*_F_ can only be estimated when *f*_P_ is measured, and only our group has done so in human PET studies with [^11^C]WAY100635. There are therefore no other datasets in which to probe these *f*_P_ effects. Setting *f*_P_ aside, group differences were observed using *BP*_P_ in (Parsey et al., 2006), but these were not replicated in (Parsey et al., 2010), while Hirvonen et al. (2008) and Meltzer et al. (2004) found differences in the opposite direction. On the other hand, no differences were found using *BP*_ND_ using the cerebellar white matter as a reference region. The aforementioned issues are restricted to *in vivo* molecular imaging, however *post mortem* autoradiography studies of the 5-HT_1A_ in MDD have also yielded mixed results, thought to be due to confounding from medication effects, comorbidities, and postmortem interval among others (Kaufman et al., 2016). In summary, this research question remains unresolved, with some review articles concluding that 5-HT_1A_R exhibits higher binding in unmedicated MDD patients relative to HV (Shrestha et al., 2012; Kaufman et al., 2016), while others conclude that binding is lower in MDD (Savitz and Drevets, 2013; Wang et al., 2016) —although the latter makes use of meta-analysis of study outcomes across different outcome measures (including indirect *BP*_ND_), reference regions, and even diagnoses, which we would assert to be an inappropriate application of meta-analysis. Besides the contrasting clinical conclusions arrived at by different surveys of the literature, the methodological questions surrounding group differences in reference quantities, and how best to contend with them has remained unclear (Parsey et al., 2010; Shrestha et al., 2012).

We have recently developed two new methods for the analysis of PET data which make use of a hierarchical multivariate modelling approach: SiMBA (Simultaneous Multifactor Bayesian Analysis) for combined quantification and statistical analysis (Matheson and Ogden, 2022), and a simplified version of this approach called PuMBA (Parameters undergoing Multivariate Bayesian Analysis) for multivariate statistical analysis following prior quantification (Matheson and Ogden, 2023). By using a hierarchical modelling strategy, SiMBA improves the accuracy of estimated parameters, and is therefore able to reliably quantify binding potential directly from rate constants without the need for a reference region since it can “borrow” information across the sample and thereby stabilize estimation of otherwise poorly identified parameters. This serves to obviate the need to rely on the imperfect cerebellar reference region to calculate these outcomes, and in simulated data we have previously shown that this approach yields outcomes with a high degree of accuracy at the individual level (Matheson and Ogden, 2022). For instance, we have previously shown in simulated [^11^C]WAY100635 data that this approach is able to reduce the error in estimates of *BP*_ND_ and non-displaceable distribution volume (*V*_ND_) by over 80% (Matheson and Ogden, 2022). Importantly, owing to the multivariate analysis strategy employed by SiMBA and PuMBA, these methods can exploit multivariate relationships among estimated pharmacokinetic (PK) parameters, which provides inferential advantages over conventional univariate analysis at the group level, including greatly improved statistical power without any corresponding increase in the false positive rate. We therefore anticipated that these methods might yield additional clarity into the study of the 5-HT_1A_R in MDD, and potentially resolve outstanding questions.

In this study, we aimed to apply multivariate analysis strategies, SiMBA and PuMBA, to study potential binding differences between MDD patients and HV, using the combined dataset from all previous studies conducted for which all outcome measures can be estimated, i.e. those which were collected at Columbia University. We applied SiMBA to study outcomes derived from a two tissue compartment model (2TCM): *BP*_ND_, *BP*_P_ and *BP*_F_, all estimated directly, in contrast to previous studies which used indirect quantification by employing a cerebellar reference region. To allow comparison, since SiMBA has not yet been extended to allow for reference tissue modelling, we use PuMBA to study indirectly-estimated *BP*_ND_ using the simplified reference tissue model (SRTM) (Lammertsma and Hume, 1996) with the cerebellar white matter as reference region.

## 2 MATERIALS AND METHODS

### 2.1 Participants and Measurement Protocols

We considered a sample of 160 individuals, including 57 HV (32 female), and 103 MDD patients (64 female). Of the patients, using the categories defined in Parsey et al. (2010), 50 were not-recently-medicated (NRM) (34 female) and 53 were antidepressant-exposed (AE) (30 female). NRM refers to having not received antidepressant medication at any point within the past four years, or being antidepressant-naive (Parsey et al., 2010). Importantly, all AE patients were either unmedicated at the time of enrollment or underwent medication washout, and none were on antidepressant medication for at least two weeks prior to PET. This sample consists of all participants from Parsey et al. (2006), Parsey et al. (2010) and Miller et al. (2013), as well as an additional 6 HV, 5 NRM patients and 12 AE patients who were not included in these studies, but with identical inclusion criteria as well as PET and MRI acquisition protocols. PET measurements were collected for 110 minutes, with 20 frames of duration: 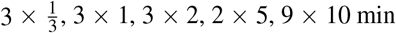.

**Table 1.**
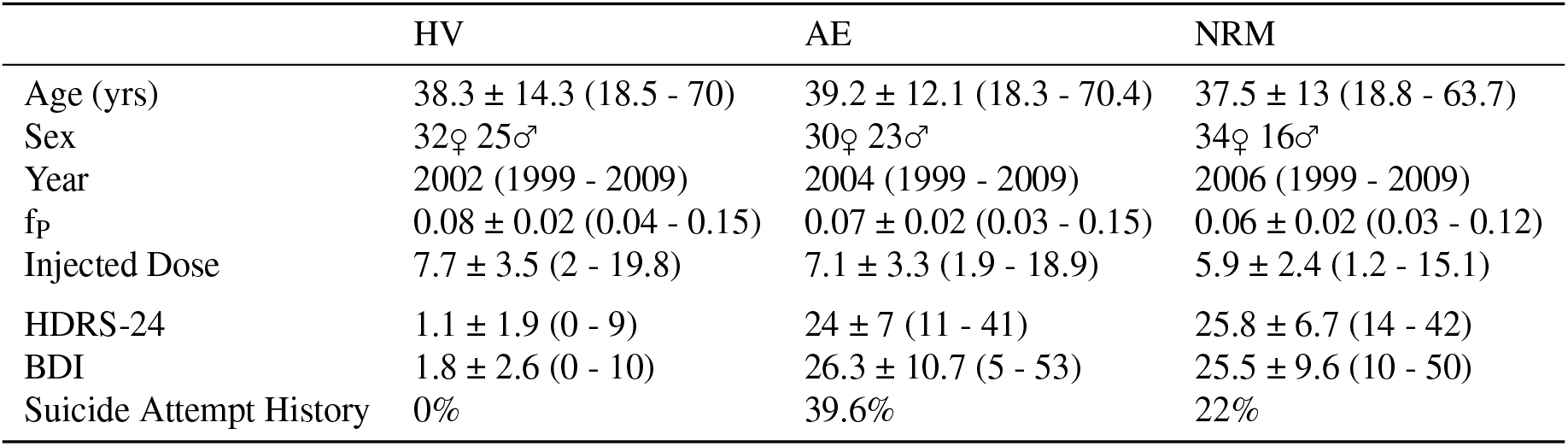
Demographic Information. Mean ± SD (Minimum - Maximum)

|  | HV | AE | NRM |
| --- | --- | --- | --- |
| Age (yrs) | 38.3 ± 14.3 (18.5 - 70) | 39.2 ± 12.1 (18.3 - 70.4) | 37.5 ± 13 (18.8 - 63.7) |
| Sex | 32♀ 25♂ | 30♀ 23♂ | 34♀ 16♂ |
| Year | 2002 (1999 - 2009) | 2004 (1999 - 2009) | 2006 (1999 - 2009) |
| $f_P$ | 0.08 ± 0.02 (0.04 - 0.15) | 0.07 ± 0.02 (0.03 - 0.15) | 0.06 ± 0.02 (0.03 - 0.12) |
| Injected Dose | 7.7 ± 3.5 (2 - 19.8) | 7.1 ± 3.3 (1.9 - 18.9) | 5.9 ± 2.4 (1.2 - 15.1) |
| HDRS-24 | 1.1 ± 1.9 (0 - 9) | 24 ± 7 (11 - 41) | 25.8 ± 6.7 (14 - 42) |
| BDI | 1.8 ± 2.6 (0 - 10) | 26.3 ± 10.7 (5 - 53) | 25.5 ± 9.6 (10 - 50) |
| Suicide Attempt History | 0% | 39.6% | 22% |

### 2.2 PET Modelling and Analysis

When applying the 2TCM (Gunn et al., 2001), we performed quantification and analysis using SiMBA (Matheson and Ogden, 2022). We defined the model with a specific binding represented using either *BP*_ND_, *BP*_P_ or *BP*_F_, estimated directly, i.e. using rate constants. Although this approach has previously been recommended against in conventional pharmacokinetic modelling of PET data using nonlinear least squares estimation (Slifstein and Laruelle, 2001) as it tends to be highly error-prone; we have previously shown that with SiMBA these outcome measures are estimated much more accurately, and with approximately 80% measurement error (Matheson and Ogden, 2022). For the *BP*_ND_ model, the PK parameters were the natural logarithms of *K*_1_, 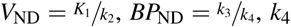, and *v*_B_. For the *BP*_P_ model, *BP*_ND_ above was replaced by 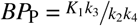. For the *BP*_F_ model, the arterial input function (AIF) was multiplied with the measured *f*_P_ value to yield an input function representing the concentration of metabolite-corrected free radioligand in arterial plasma. Hence, the parameters of the latter model were the natural logarithms of 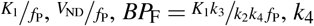, and *v*_B_. In all models described, the *v*_B_ parameter represents the blood volume fraction, i.e. the fraction of the regional volume comprised of blood vasculature and whose time activity curve (TAC) is represented by the measured whole blood curve. Because whole blood was not measured in this study, we made use of whole plasma as a proxy for whole blood, as performed in previous studies using these data (Parsey et al., 2006, 2010; Miller et al., 2013).

When applying SRTM (Lammertsma and Hume, 1996), we were not able to use SiMBA since the reference region-based functionality has not yet been developed. Rather, we made use of PuMBA (Matheson and Ogden, 2023), using the parameters estimated using conventional nonlinear least squares (NLS) estimation, with the cerebellar white matter as reference region. NLS estimation was applied using `kinfitr` (Matheson, 2019; Tjerkaski et al., 2020). For NLS modelling, weights were defined according to the default `kinfitr` method, i.e. the square root of the product of the frame duration and the mean TAC value, after being un-corrected for radioactive decay. As input parameters for the PuMBA model, we used the natural logarithms of *R*_1_, 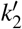 and *BP*_ND_.

Univariate analysis was performed using linear mixed effects (LME) modelling, in which the natural logarithm of the relevant binding potential (*BP*) outcome was defined as the dependent variable, with covariates for region, sex, age, a group-by-region interaction term, as well as a random intercept for individual. SiMBA and PuMBA analyses were defined with the same set of covariates for *BP*, but these models additionally require defining regression models for each of the covariates. We defined partial pooled (random effect) deviations for region, individual and TAC, as previously described (Matheson and Ogden, 2023, 2022). For the 2TCM, age and sex were defined as fixed effect covariates for the natural logarithm of *K*_1_, and for *V*_ND_. For PuMBA analysis of the SRTM results, age and sex were defined as fixed effect covariates for the natural logarithm of 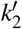.

For the Bayesian multivariate models, we used moderately informative priors for the global parameter intercepts, as described previously (Matheson and Ogden, 2023, 2022), and weakly informative regularising priors otherwise. Zero-centred regularising priors were defined over the standard deviation of the partially pooled effects, with standard deviations defined in such a way as to exclude implausible magnitudes of variation. For the differences between MDD patients and HV, we defined zero-centred regularising priors with a normal distribution centred at zero with a standard deviation of 0.2. This implies that our prior beliefs are that no group differences are most likely, and that we are progressively more skeptical of differences of larger magnitude. The prior assigns 18% of its prior probability to differences less than 10%, 64% to differences less than 20%, and 81% to differences less than 30%. The prior is two-sided, meaning that our model assigns only 9% prior probability to increases greater than 30%, and 9% to decreases greater than 30%. The priors are more fully described in Supplementary Materials S1.

All modelling and analysis were performed using `R 4.0.5 (Shake and Throw)` (R Core Team, 2022). Linear mixed effects model analysis was performed using `lme4` (Bates et al., 2015). Markov Chain Monte Carlo fitting was performed using `STAN` (Carpenter et al., 2017) together with the `brms` R package (Bürkner, 2017).

## 3 RESULTS

### 3.1 Group Comparisons

#### 3.1.1 Plasma Free Fraction

We observed group differences in the natural logarithm of *f*_P_ (*F*_2,157_ = 10.1, p *<* 0.001) (Fig 1A), with differences between HV and both AE patients (Estimated difference in *f*_P_ = 22%, p = 0.003) and NRM patients (Estimated difference in *f*_P_ = 28%, p *<* 0.001), but no significant differences between the two patient groups (Estimated difference in *f*_P_ = 6%, p = 0.631). If these differences in measured *f*_P_ values reflect true differences in the plasma free fraction, then it is important that they be accounted for during quantification by estimating *BP*_F_. However, due to the inhomogeneous distribution of sampling from the different participant groups over the years of data collection, e.g. that the majority of HV were imaged before the patient groups, it is also possible that they could be explained by experimental drift in the measurement of *f*_P_. If this is the case, the *BP*_F_ estimates will be systematically biased, and global group differences will be artificially induced.

**Figure 1.**
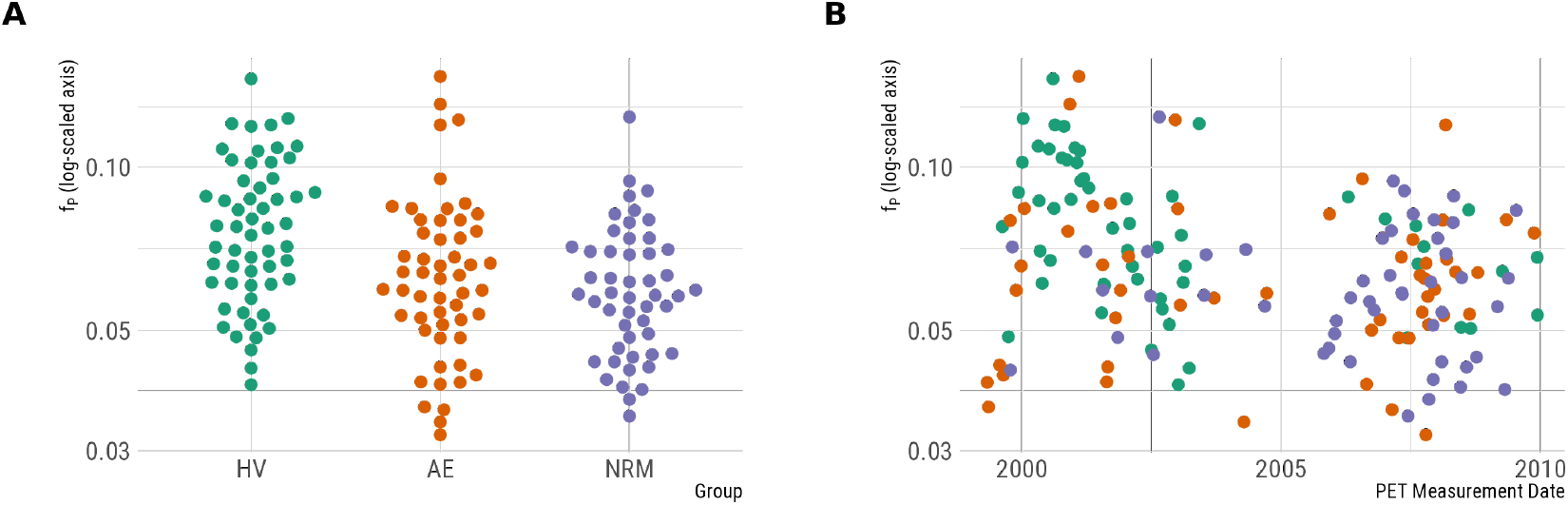
Potential explanatory factors for measured plasma free fraction values. A. Group differences in the plasma free fraction. B. Plasma free fraction as a function of the date of PET measurement.

To explore this possibility, we fit an additional analysis with fixed effects for group differences as before, but including a smooth term for the date of PET measurement to describe any potential experimental drift. The model was fitted using the `mgcv` (Wood, 2011) package using a thin-plate spline regression with 10 degrees of freedom with the smoothing penalty estimated using restricted maximum likelihood estimation. Using this model, the smooth experimental drift term was statistically significant (*F*_8.1_ = 5.00, p *<* 0.001), while the group differences were no longer significant (*F*_2_ = 1.225, p = 0.297). This suggests that it is at least plausible that the group differences in *f*_P_ might be caused by experimental drift, although neither possibility can be conclusively ruled out based on the data at hand because of the inhomogeneous time distribution of the groups.

#### 3.1.2 Univariate Analysis

Using conventional univariate LME analysis of PET outcomes, we broadly replicate the results of previous studies (Parsey et al., 2006, 2010; Miller et al., 2013) and their associated discrepancies between the results obtained using different outcome measures (Fig 2, A-C). In contrast to previous analyses, all 2TCM outcomes are estimated directly from rate constants, as opposed to indirectly using the cerebellum as a reference region, in order to be comparable to the multivariate analyses. Also, in contrast to previous analyses, the ratio of *K*_1_/*k*_2_ in each region is not constrained to be equal to the same ratio in the white matter cerebellum (i.e., a constrained 2TCM was used in previous analyses).

**Figure 2.**
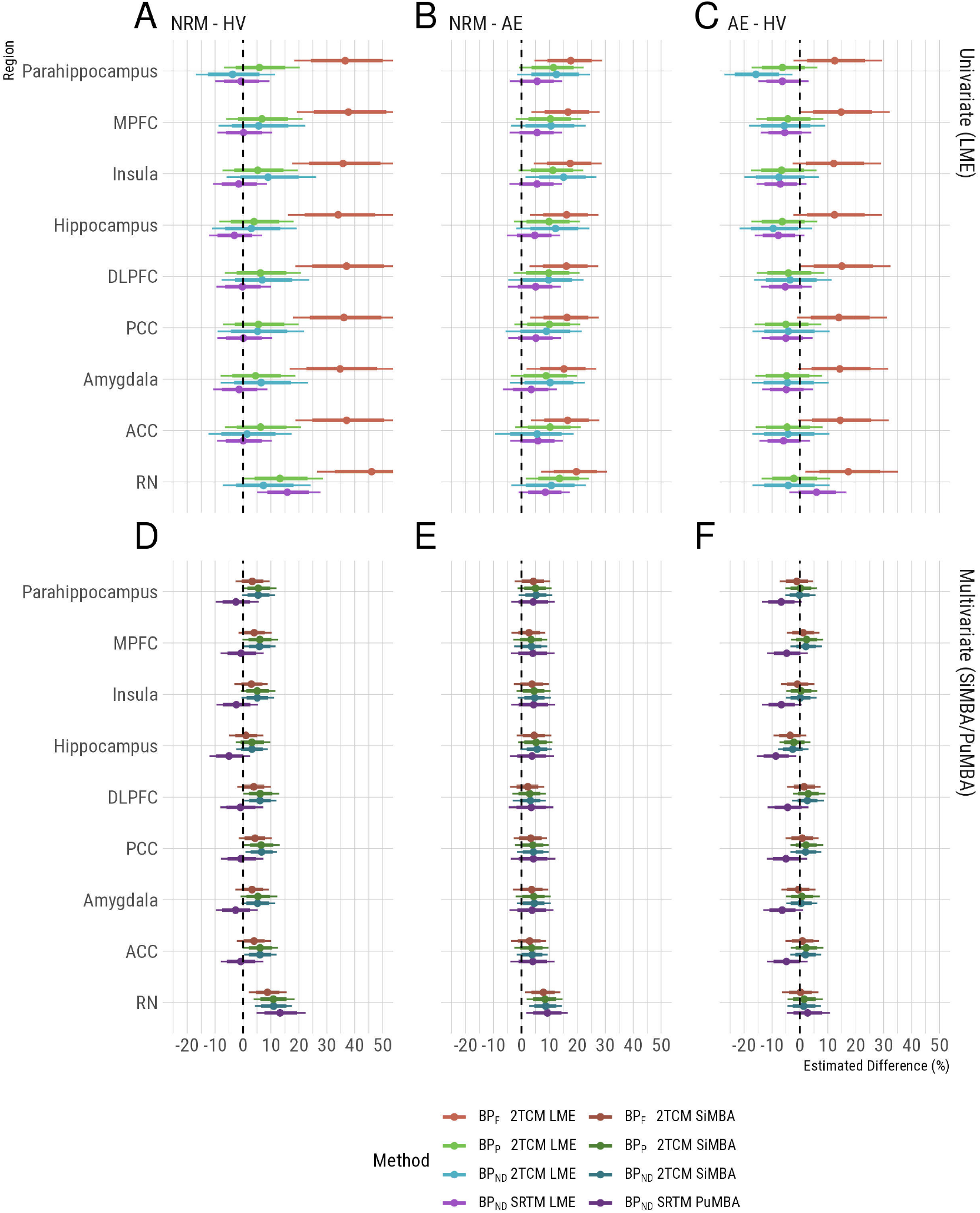
Group differences and their associated uncertainty, presented as percentage differences between groups. Colors represent the outcome measure used, in which lighter colors represent univariate LME results and darker colors represent multivariate results. Uncertainty intervals represent the 80% and 95% confidence/credible intervals with the thicker and thinner bands respectively. Regional abbreviations are as follows: DLPFC is dorsolateral prefrontal cortex, MPFC is medial prefrontal cortex, ACC is anterior cingulate cortex, PCC is posterior cingulate cortex, and RN is the raphe nuclei

Focusing on the differences between NRM MDD patients and HV (Fig 2, A), which has been the primary focus of previous studies, large global differences are observed between groups using *BP*_F_. However, owing to the large differences in *f*_P_ (Fig 1A) which may or may not be caused by true biological differences, it is unclear whether these differences in *BP*_F_ are genuine or a methodological artefact. The results for *BP*_P_ and *BP*_ND_, in which *f*_P_ differences are not corrected for, do not exhibit the same global differences between groups (Fig 2, A). In previous studies of these data (Parsey et al., 2006, 2010; Miller et al., 2013), these outcomes have been estimated indirectly, i.e. using both arterial plasma as well as a reference tissue, due to concerns that direct estimation may compromise the accuracy of estimated outcomes (Slifstein and Laruelle, 2001). However the fact that we fully replicated the outcome measure discrepancies observed in previous studies of these data suggests that our use of direct estimation is unlikely to be a major source of differences between this and previous analyses.

Lastly, using indirect estimation of *BP*_ND_ using SRTM with the cerebellar white matter as a reference region, we observe higher *BP*_ND_ in NRM patients relative to HV in the raphe nuclei (RN).

#### 3.1.3 Multivariate Analysis

Using the multivariate analysis approaches, SiMBA and PuMBA, the uncertainty intervals are much narrower than those calculated using the univariate methods as shown previously (Matheson and Ogden, 2022) and, importantly, the estimates are broadly consistent across all four outcomes (Fig 2, D-F). Focusing on the differences between NRM patients and HV again (Fig 2, D), all outcomes show regionally-specific higher binding potential in the RN of NRM patients relative to HV. NRM patients also show higher RN binding potential compared to AE patients consistently across outcome measures (Fig 2, E). These similarities between outcomes are observed in spite of the differences in *f*_P_ between groups which would affect the estimation of *BP*_F_ but not the other outcome measures. Similarly, SRTM *BP*_ND_ also exhibits a similar pattern of results as the 2TCM outcomes above (Fig 2, D-F), in spite of both being an indirect estimate of *BP*_ND_, i.e. estimated without the use of arterial blood data; as well as being estimated using multivariate analysis of a different set of PK parameters, i.e. those estimated by SRTM.

In all the remaining regions where post-synaptic heteroreceptors are located, there was a tendency for slightly higher binding in NRM patients compared with both HVs and AE patients for the 2TCM outcomes, however these estimates had a magnitude of approximately half of that observed in the RN, and mostly had 95% credible intervals which overlapped with 0. In order to study the regional specificity further, we fit additional models for each outcome measure with fixed effects for overall group differences in the RN and for the serotonin projection regions, with random slopes for the serotonin projection region (i.e. allowing variation in regional differences around their overall mean). Comparing the magnitude of group differences in the RN with the mean difference in projection regions, we show that differences are larger in the RN 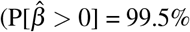 for 2TCM *BP*_F_, 98.9% for 2TCM *BP*_P_, 99.3% for 2TCM *BP*_ND_ and 99.5% for SRTM *BP*_ND_) comparing NRM patients with controls. However, BP is still higher on average in projection regions in NRM patients compared to controls for 2TC models (although the 95% credible interval partially overlaps with 0 for *BP*_F_): 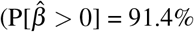 for *BP*_F_, 97.7% for *BP*_P_, and 98.5% for *BP*_ND_). Within the projection regions, there is minimal regional variation in the magnitude of differences, i.e. the random slopes: the standard deviation of ΔlogBP ¡ 0.01 for all outcome measures for both NRM - control comparisons as well as AE - control comparisons. All estimates and directional probabilities comparing regional differences as both fixed effects and random slopes are provided in Supplementary Materials S3.

Notably, the pattern of results for the projection regions was inconsistent between direct (2TCM) and indirect (SRTM) quantification approaches. For the NRM - HV comparisons, all direct approaches demonstrate elevations in binding potential in NRM patients compared with HV for projection regions, while the indirect SRTM approach does not 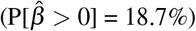. Similarly, for the AE - HV comparisons (Fig 2, F), all direct quantification approaches are consistent with no differences, yet the SRTM indirect quantification indicates lower binding potential in AE patients in projection regions 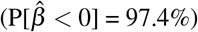. One possible explanation for these inconsistencies could be that cerebellar white matter binding is higher in MDD compared with HV. While Parsey et al. (2010) showed that *V*_T_ in cerebellar grey matter was elevated in MDD groups compared to HV, they did not observe statistically significant differences in cerebellar white matter. To explore this possibility, we fit an additional exploratory SiMBA model examining group differences in *BP*_F_ where cerebellar grey and white matter regions were additionally included as regions of interest. This model indicated that *BP*_F_ was higher in MDD, regardless of antidepressant exposure, compared with HV in both cerebellar grey matter (AE - HV: 22% [12 - 32%] 95% CI, 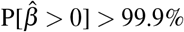; NRM - HV: 39% [28 - 50%] 95% CI, 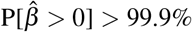) and cerebellar white matter (AE - HV: 9% [0.2 - 20%] 95% CI, 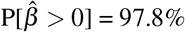; NRM - HV: 13% [4 - 23%] 95% CI, 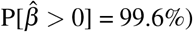). This finding supports the hypothesis proposed by Parsey et al. (2010) that the small degree of specific binding in cerebellar grey matter is greater in MDD; and that this may also be the case for cerebellar white matter, which might therefore explain the bias observed in the SRTM results above.

Lastly, we did not observe the associations between sex and binding which have been reported in prior studies (Parsey et al., 2006, 2010). However, we did observe sex differences in *K*_1_, with males showing a lower rate of transfer from arterial plasma to tissue compared with females. This outcome was reasonably consistent across all 2TC models examining *BP*_F_ (−5.1%, 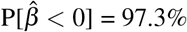), *BP*_P_ (−5.6%, 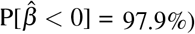) and *BP*_ND_ (−5.9%, 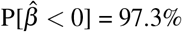).

### 3.2 Individual-Level Estimate Associations

Given the consistency of the group-level inferences between outcome measures despite group differences in *f*_P_, we sought to compare individual-level outcomes. To this end, we extracted the partially pooled individual-level estimates (i.e. 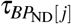 in Matheson and Ogden (2022)), also known as the random effects, for the binding potential outcomes. These figures represent the mean individual binding potential values after adjustment for all of the included covariates (including group membership) as well as regional differences. This allows us to examine the consistency of individual binding potential estimates across outcomes. We extracted the individual-level binding potential estimates for SiMBA with *BP*_F_, SiMBA with *BP*_ND_, and for PuMBA with SRTM *BP*_ND_.

Comparing direct estimates, we observe a high degree of correspondence (r ¿ 0.99) between *BP*_P_ values estimated by the 2TCM with and without correction for *f*_P_ (Fig 3A). In contrast, we observe weaker associations (r = 0.59) between *BP*_ND_ values estimated by these two models (Fig 3B). Keeping in mind that *BP*_P_ is equal to the product of *BP*_ND_ and *V*_ND_, this implies that correction of the AIF for *f*_P_ differences alters the balance of *BP*_ND_ and *V*_ND_ estimates at the individual level, but that changes in each one of these parameters are accommodated for by corresponding changes in the other within the SiMBA model. When instead examining the correspondence between direct (arterial-blood based 2TCM) and indirect (reference-region-based SRTM) estimates of *BP*_ND_, we see little to no association between estimates either with (r = 0.23, Fig 3C) or without (r = 0.19, Fig 3D) correction for *f*_P_.

**Figure 3.**
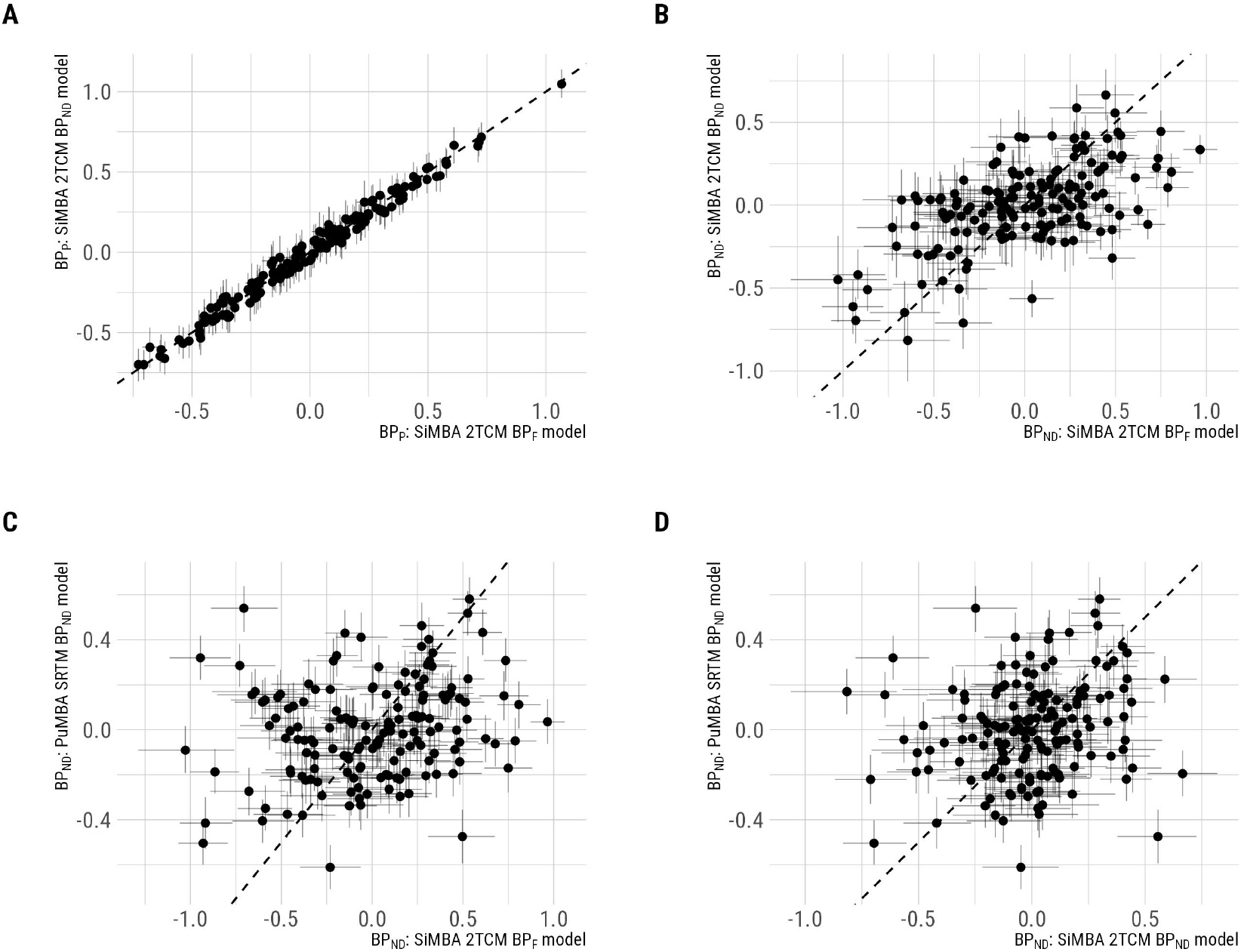
Individual-level binding potential estimates compared between models, and lines of identity. A. Comparisons of *BP*_P_ estimated by SiMBA with and without the correction of the AIF for *f*_P_. B. Comparisons of *BP*_ND_ estimated by SiMBA with and without the correction of the AIF for *f*_P_. C. Comparison of *BP*_ND_ estimated by SiMBA with correction of the AIF for *f*_P_ and indirect estimates calculated using PuMBA with SRTM. D. Comparison of *BP*_ND_ estimated by SiMBA without correction of the AIF for *f*_P_ and indirect estimates calculated using PuMBA with SRTM.

### 3.3 Correlation Matrices

Both SiMBA and PuMBA models exploit the multivariate associations between estimated PK parameters to improve inferential power. One of the assumptions of these models is that the (random effect) correlations between PK parameters are shared across the total sample after correction, and that they do not differ between groups. The model is defined in such a way that it should be robust to small deviations from this assumption, as this residual variance can be accommodated by Individual × Region (i.e. TAC) variance; however large deviations between groups could potentially be problematic. To test this assumption, we fit a SiMBA *BP*_ND_ model to each of the three subgroups independently, and extracted the estimated individual-level correlation matrices for comparison. We selected the SiMBA *BP*_ND_ model, as *BP*_ND_ is less stable than *BP*_P_, and ought to exaggerate any potential differences between groups, especially considering the relatively smaller sample sizes of each separate group. The results are shown in Figure 4. Based on the high degree of similarity of these matrices, we do not anticipate that this is a source of appreciable bias in the results of our models.

**Figure 4.**
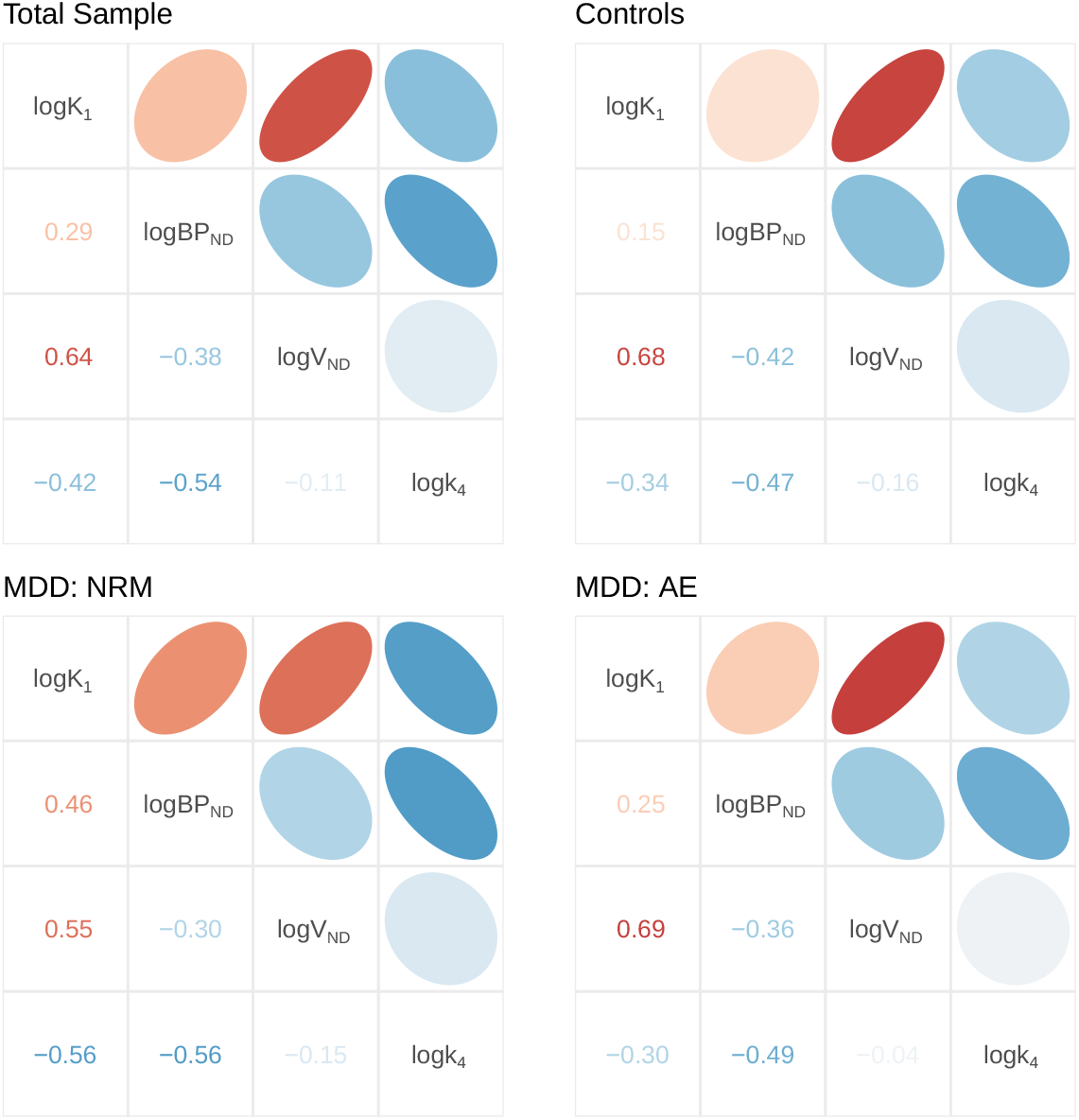
Individual-level random effect correlation matrices were estimated for the total sample as well as each patient subgroup independently. The similarity of these matrices suggests both that their estimation is robust, but also that there do not appear to be any clear systematic differences between groups that are not more likely to be explained by estimation inaccuracies.

## 4 DISCUSSION

We have applied novel hierarchical multivariate analysis methods, that we have previously shown to improve quantitative accuracy and inferential efficiency (Matheson and Ogden, 2022, 2023), to the largest-to-date sample of [^11^C]WAY100635 PET data, including full arterial sampling and measurement of the radioligand *f*_P_, to study differences between HV and participants with MDD. In contrast to previous studies, using these new methodological approaches we find highly consistent results using *BP*_F_, *BP*_P_ and *BP*_ND_, and using both the 2TCM and SRTM pharmacokinetic models. Concordance across binding outcome measures and models is expected in theory, but has not been observed in previous studies employing traditional PET analysis methods to study the 5-HT_1A_ receptor in MDD. Moreover, while previous studies found generalised increases across the brain in not-recently-medicated MDD patients relative to controls, which were restricted to *BP*_F_, our current study is novel in that it both corroborates these findings across all outcome measures, as well as extending these findings by showing that these differences are most pronounced in the RN. This regional specificity, however, is consistent with the particular neurobiological role played by 5-HT_1A_ autoreceptors in the RN relative to heteroreceptors in the the rest of the brain. This study may therefore help resolve the longstanding discrepancies observed in studies employing PET imaging to quantify the 5-HT_1A_R binding in MDD.

The regional specificity of our findings is of particular interest given the particular role of the 5-HT_1A_R in the RN, where it functions as a somatodendritic autoreceptor, as opposed to its role as a heteroreceptor in the rest of the brain. These autoreceptors control serotonin neuron firing and serotonin release throughout the brain through hyperpolarizing the serotonin neurons via a potassium channel in response to increased extracellular serotonin concentrations (Barnes and Sharp, 1999; Hoyer et al., 2002; Montalbano et al., 2015). It has long been speculated based on rodent studies (Blier and De Montigny, 1983; Gardier et al., 1996; Piñeyro and Blier, 1999) that 5-HT_1A_ autoreceptors mediate the delayed onset of action of drugs which increase serotonin levels, such as selective serotonin reuptake inhibitors (SSRIs). Initially, increasing extracellular serotonin concentrations results in 5-HT_1A_ autoreceptor-induced hyperpolarization of serotonin neurons, and inhibition of firing and serotonin release. With prolonged treatment, these receptors become desensitised and eventually downregulated through internalization, resulting in enhanced firing and serotonin release and emergence of an antidepressant effect (Blier and De Montigny, 1983; Gardier et al., 1996; Piñeyro and Blier, 1999; Richardson-Jones et al., 2010), leading to a gradual increase in treatment effects over several weeks (Taylor et al., 2006). In mice genetically engineered to specifically express high or low levels of 5-HT_1A_ autoreceptors, the mice with low levels show less depression-like behavior, while also exhibiting greater antidepressant response to SSRI medication (Richardson-Jones et al., 2010). Our findings are consistent with these results: we observe higher 5-HT_1A_ autoreceptor binding in not-recently-medicated patients compared to both HV, suggesting the possibility of its role in the pathophysiology of MDD, as well as compared to antidepressant-exposed MDD patients, suggesting the possibility of residual treatment effects downregulating 5-HT_1A_R binding in the AE group as shown in Gray et al. (2013).

The question remains why this pattern of results was not observed in previous studies: if binding potential for [^11^C]WAY100635 in the RN is higher in unmedicated MDD, then why have results been so inconsistent? For *BP*_P_ in particular, results have been inconclusive, despite its being less sensitive to radiometabolite contamination than *BP*_ND_, and less sensitive to any potential methodological biases in the measurement of *f*_P_ than *BP*_F_. We offer two potential answers to this question. Firstly, these differences were observed in the RN: this region is very small and its TACs have higher measurement error compared to the other brain regions examined here. For this reason, its estimation is more susceptible to quantification inaccuracies than the other regions. SiMBA and PuMBA both make use of hierarchical modelling across both individuals and brain regions, which allows them to effectively borrow strength from the rest of the dataset to improve inferences made. For SiMBA in particular, quantification is performed within the model, and using the rest of the data, resulting in substantial improvements to the accuracy of quantification at the individual level - with greatest improvements observed for the RN (Matheson and Ogden, 2022). Secondly, previous studies have made use of univariate analysis methods to study this research question. This means that binding potential values are estimated for each region of each individual and entered into a statistical test, while all the other PK parameters estimated by the model are discarded. We have shown for both SiMBA (Matheson and Ogden, 2022) and PuMBA (Matheson and Ogden, 2023) that there is additional information within these discarded parameters and their covariance structures, which can be exploited to improve, sometimes dramatically, the power of group-level inferences made using binding potential —particularly with larger sample sizes. In simulations of [^11^C]WAY100635 data, using *BP*_P_ as an outcome measure, multivariate inferential power was equivalent to that of a sample approximately twice the size using univariate analysis (Matheson and Ogden, 2022). To this end, as part of this study, we performed a power analysis to estimate sample size required to test the differences observed in this study in *BP*_P_ between NRM patients and HV in the RN for *BP*_P_ using a univariate *t*-test. The results of the power analysis are reported in Figure 5, showing that a sample size of 158 individuals (95% CI: 57 - 1176) would be required in *each group* to achieve 80% power with conventional univariate *t*-test, owing to the small group differences observed which correspond to a Cohen’s *d* of 0.32 (95% CI: 0.11 - 0.53). To contextualise the magnitude of these differences, this figure implies that selecting a healthy control and a NRM MDD patient at random, the probability of the NRM patient having a higher binding potential than the control is only 58.7% (95% CI: 54.4% - 64.6%) due to the substantial degree of overlap between groups (Supplementary Materials S4). Hence, even including all participants from all previous studies of this research question (Shrestha et al., 2012; Wang et al., 2016) in a single *t*-test, even after correcting for centre differences, would be underpowered to detect such a difference. As such, while these differences may be informative from a scientific perspective, their clinical utility is likely limited.

**Figure 5.**
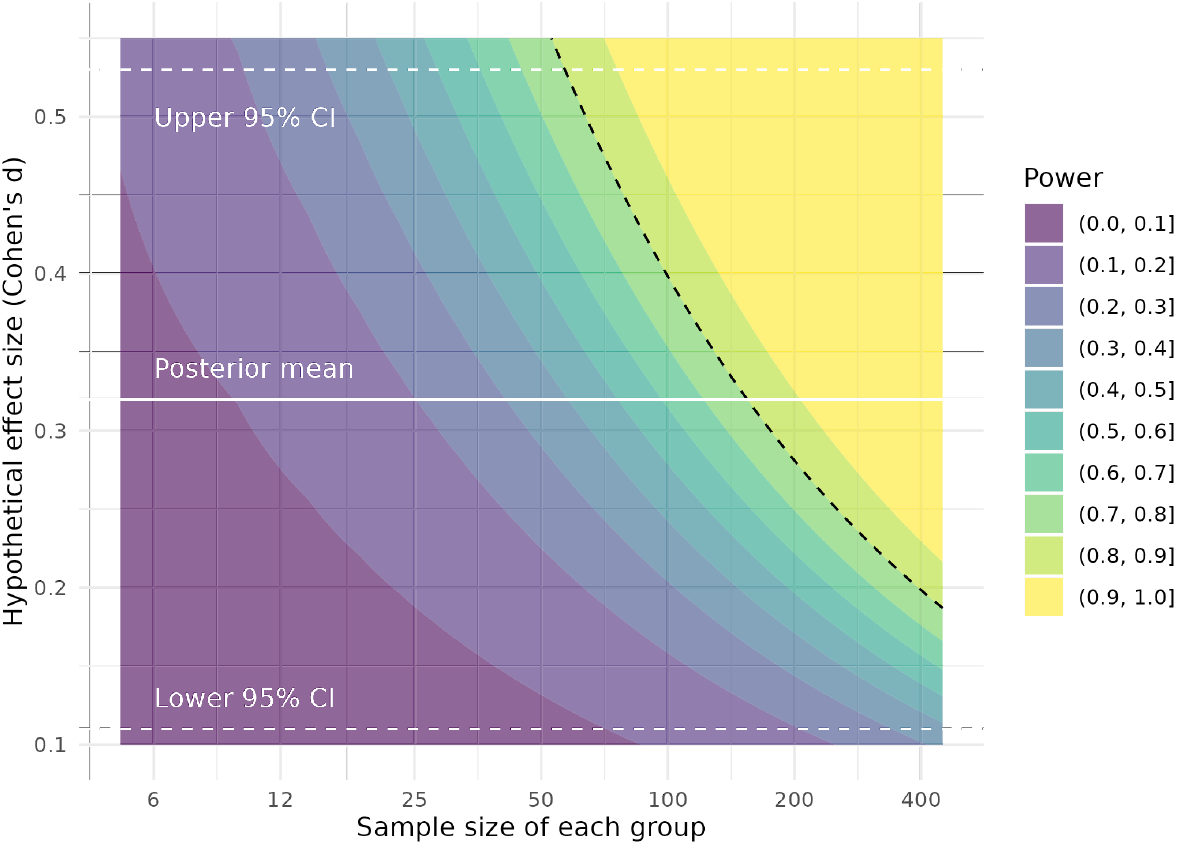
Based on the results of our analysis, the required samples sizes for a univariate *t*-test comparing BP_ND_ between HV and not-recently medicated MDD are shown as a function of the hypothetical true effect size. The dotted black line shows where 80% power lies.

One surprising outcome was the overall similarity of the group-level inferences across the different binding potential outcomes in spite of the dissimilarity of the individual-level binding potential estimates, also known as the random effect estimates. This shows that the hierarchical parameters and their shrinkage were different for each of these outcome parameters, mirroring the variability of previous results across different outcome parameters. However, by acknowledging and exploiting the multivariate covariance structure in the data between all PK parameters simultaneously, our models were able to make inferences which were more resilient to these differences. This raises the important issue of how stable these multivariate relationships really are: as it is implemented, the model requires that the correlation structure is reasonably consistent between groups. To this end, by fitting the model to each subgroup of the data independently and showing that the estimated correlation structure was highly similar between groups, we conclude that this is unlikely to have been a cause for concern.

The most clear differences in the results were between outcomes using direct and indirect means. Not only were there differences in group-level inferences, but the individual-level *BP*_ND_ estimates were also uncorrelated. The problems with indirect estimation of *BP*_ND_ for [^11^C]WAY100635 have been described previously (Osman et al., 1998; Parsey et al., 2005, 2010): owing to the low distribution volume of the cerebellar grey and white matter (and the low *V*_ND_ of [^11^C]WAY100635 more generally), the small absolute deviations in the distribution volume caused by contamination by small concentrations of radioactive metabolites constitute large proportional deviations. This therefore exerts an outsized influence on *BP*_ND_ compared to, for instance, *BP*_P_. However, the use of indirect estimation of *BP*_ND_ for [^11^C]WAY100635 with SRTM are not limited to this issue. The SRTM model makes four primary assumptions (Lammertsma and Hume, 1996; Salinas et al., 2014). Firstly, the model assumes that there should be no displaceable component in the reference region. This is, however, not true of the cerebellar grey matter (Parsey et al., 2005; Hirvonen et al., 2007; Parsey et al., 2010), and potentially of the white matter to a smaller extent due to spill-in from partial volume effects. Secondly, SRTM assumes that the blood volume to the tissues is negligible: given that uptake is so low in the cerebellum, this assumption is unlikely to be met. Thirdly, SRTM assumes that there are one-tissue compartment model (1TCM) kinetics in both the target and reference regions, which is not met, as the 1TCM does not provide an adequate fit in either. Lastly, SRTM makes the assumption that *V*_ND_ is the same in all regions. Using SiMBA, however, the model evaluates variation in regional mean *V*_ND_ estimates (i.e. 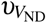) for the 2TC models (Matheson and Ogden, 2022). From the model including the cerebellar regions, *V*_ND_ is estimated to be lowest compared to the mean in the cerebellar white matter (−35% 95% CI: -42.6% - -28.1%), followed by the raphe nuclei (−16.5% 95% CI: -29.9% - -6.1%) and the cerebellar grey matter (−11.7% 95% CI: -23.5% - -1.3%), while all the other regions had credible intervals which overlapped with the mean value. This provides evidence that this assumption may also not be fulfilled. In sum, there is reason to believe that none of the four primary assumptions of SRTM are fulfilled for [^11^C]WAY100635. For this reason, as well as those discussed in Shrestha et al. (2012), we consider the outcomes obtained using 2TCM to be more reliable than those of SRTM, however the consistency of its inferences to those of the 2TCM showing most pronounced differences in the RN is interesting nonetheless. Similarly, we stress that retrospective analyses which treat results obtained using reference tissue modelling, particularly using cerebellar grey matter as a reference region, as equivalent to those obtained using the 2TCM should be interpreted with caution, and recognising the inherent limitations of this outcome and its sources of bias for [^11^C]WAY100635 PET.

Unfortunately, most of the studies of MDD using [^11^C]WAY100635 have been performed using reference tissue modelling. This is because this approach does not require arterial sampling, which is costly, technically complex, as well as uncomfortable for participants. Aside from the data included in this study, there exist only two other published PET studies examining [^11^C]WAY100635 in MDD in which arterial plasma data was collected: Hirvonen et al. (2008) included 21 patients, and Meltzer et al. (2004) included 17 patients. Neither of these two studies, however, collected *f*_P_ data which would have allowed calculation of *BP*_F_. Hence, the data included in this study constitutes the entirety of PET data examining MDD with [^11^C]WAY100635 for which all outcome measures can be estimated. This was discussed in Shrestha et al. (2012) which concludes, owing to the *f*_P_ differences observed in Parsey et al. (2006) and Parsey et al. (2010), that *BP*_F_ is the only outcome measure which can be used to directly study this question because, if there are true differences in *f*_P_ between groups, then any analysis examining *BP*_P_ in which these differences are not corrected for will provide biased results. This leaves *BP*_ND_, but as discussed earlier, this outcome measure is compromised using reference tissue modelling, and its estimation using the 2TCM is considered too unreliable using conventional methods. For this reason, the present analysis is performed using the sum total of the data for which it is possible to examine these outcome measure discrepancies even though they were all collected at only one PET centre. However, given the similarities of outcomes across outcome measures observed here, this implies that reanalysis of the other datasets for which arterial sampling was performed might also be valuable —even if *f*_P_ data was not collected.

Notably, despite the large, and statistically significant, group differences in *f*_P_, its influence on our inferences about group differences in binding potential appears to have been minimal using SiMBA. From Figure 3A, we can see that although individual estimates of *BP*_P_ in SiMBA models are highly correlated with- and without correction of the AIF by *f*_P_, they are not identical. This is because the SiMBA model performs hierarchical shrinkage, or regularization, of parameter estimates resulting in slightly different estimates when *f*_P_ is incorporated into the model. In contrast, using conventional modelling with no shrinkage, the estimates would have been perfectly associated with one another. Because SiMBA makes use of *multivariate* shrinkage in particular, it is able to make use of the covariance structure between all of the estimated PK parameters to constrain its estimation more effectively compared with univariate shrinkage. The fact that the group differences in *f*_P_ waned in the context of the multivariate hierarchical model does provide indirect support for the hypothesis that these differences may, in part, be caused by methodological drift as opposed to true biological differences. This would have been especially problematic due to the control groups being mostly overlapping between Parsey et al. (2010), Parsey et al. (2006) and Miller et al. (2013). This speaks to the importance of recruiting patient and control groups concurrently and homogeneously over time to avoid the potential for such complications, where it becomes difficult to disentangle biological factors from methodological drift.

The present results might be interpreted to imply that correction for *f*_P_ is unnecessary, however we would not espouse this position. As discussed above, based on the results of this data alone, we cannot know whether the observed group differences are true or due to methodological factors. The reason for adjusting the AIF for *f*_P_ is that radioligands, like drugs, bind to plasma proteins, establishing a binding equilibrium, and that only the free fraction of the radioligand not bound to proteins can cross the blood-brain barrier to reach the target of interest. If there are true differences in *f*_P_ between groups caused by some biological difference, then it is essential that these be corrected for.

Measurement of *f*_P_ is most commonly performed using one or more arterial plasma samples extracted from the subject prior to radioligand injection, and adding radiolabelled compound before assaying the protein-free fraction using ultrafiltration (Gandelman et al., 1994). In this study, we incorporated measured *f*_P_ values into our SiMBA model as a scalar quantity, by adjusting the AIF accordingly. While this is consistent with the conventional approach of correcting the estimated *BP*_P_ value by a scalar *f*_P_ value, it does not allow for any potential *in vivo* dynamics of *f*_P_ over time. Using ultrafiltration, as was performed in this study, it is not possible to measure changes in blood *f*_P_ over the first minutes of the scan due to its long analysis time. On the other hand, it has been reported that equilibrium concentrations of plasma protein binding may even be reached within milliseconds (Peletier et al., 2009; Smith et al., 2010), in which case *f*_P_ can be considered effectively stable. We therefore believe that this approach constitutes a sufficiently good approximation for the data-generating process in order to accommodate differences in *f*_P_ values at equilibrium; which is indirectly supported by the similarity of our inferences between outcome measures.

In summary, in this study we show that multivariate analysis methods may resolve the longstanding debate over the results of studies of [^11^C]WAY100635 in MDD, and suggest that NRM patients have higher 5-HT_1A_R binding in the RN compared to both HV and AE patients - although these differences are rather small. We propose that these methods have broad applicability for answering other research questions for which *in vivo* PET imaging is used to study psychiatric, neurological or even pharmacological questions.

## 5 DATA AND CODE AVAILABILITY

The analysis code used here is based on the R and STAN shared previously from our previous methodological paper presenting this approach (https://github.com/mathesong/SiMBA_Materials) and applied to new data. The specific code used to analyse the current data can be shared upon reasonable request after removal of sensitive patient information. The majority of the current dataset is re-analysed from previous studies, including all participants from Parsey et al. (2006), Parsey et al. (2010) and Miller et al. (2013). An additional 23 participants were also collected in the same data collections, but who were not included in the previous studies.

## 6 ETHICS

All subjects gave written informed consent prior to participation, and all studies were approved by the regional ethics committees.

## 7 AUTHOR CONTRIBUTIONS

GJM, JMM, JJM and RTO conceived and designed the analysis. FZ and EAB helped with retrieval, quality control and harmonisation of data from the different sources. GJM and RTO performed the analysis, with domain expertise and suggestions from JMM and JJM. GJM and RTO wrote the first draft of the paper, and all authors provided suggestions and revisions for the final draft.

## 9 ACKNOWLEDGMENTS

We would like to thank Dr Ramin Parsey for his thoughtful feedback and suggestions during the preparation of this manuscript, as well as the comments and feedback from the MIND research group at NYSPI. This work was supported by Hjärnfonden (PS2020-0016), Vetenskapsrådet (2020-06356), NIH grants P50MH090964 and R01EB024526.

## 10 SUPPLEMENTARY MATERIALS

### 10.1 Supplementary Materials S1: PET Quantification Information

PET quantification involves fitting pharmacokinetic (PK) models to the time course of measured radioactivity concentrations in the brain, termed time activity curves (TACs). Using these models, PET allows for quantification of various binding potentials (*BP*), i.e. the equilibrium concentration of radioligand specifically bound to the target protein, normalized to a reference concentration. While *in vitro* studies can measure the concentration of the target protein available to bind the radioligand, *B*_avail_, this is not possible from a single *in vivo* PET examination. Rather, PET researchers make use of binding potential as a measure of specific binding, which is equal to 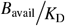, where *K*_D_ represents the affinity of the target for the radioligand. Hence, as long as *K*_D_ is unchanged, then *BP* should be proportional to *B*_avail_ and can be used as an effective surrogate. However, in order to calculate binding potential, we do so using a suitable reference quantity. As such, there are several different types of binding potential which differ from one another as a function of which reference quantity is used for their calculation, i.e. *BP*_X_ where X describes the reference quantity.

*BP*_F_ represents the gold standard PET outcome measure which is, in the absence of active transport, equal to 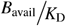. The estimation of *BP*_F_, however, requires measurement both of the metabolite-corrected radioactivity in arterial plasma during the PET measurement, as well as the protein-free fraction of the radioligand in arterial plasma, *f*_P_ (Innis et al., 2007; Parsey et al., 2010). Measurement of *f*_P_ tends to be notoriously prone to measurement error due both to estimation inaccuracy, and especially so when *f*_P_ values are low as is the case for [^11^C]WAY100635. Moreover, measurement of *f*_P_ tends to be sensitive to small experimental biases such as the temperature or pH of plasma samples, the latter of which can even vary as a result of storage time (Hinderling and Hartmann, 2005; Kochansky et al., 2008). Hence, even if data show a reasonably high degree of consistency within individuals, measured *f*_P_ values could feasibly be impacted by unexpected experimental confounds over time. Measured values can therefore suffer from both variance and bias. On the other hand, *BP*_P_ which is equal to 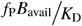, does not require the measurement of *f*_P_ - however it is sensitive to differences in plasma protein binding between individuals. For this reason, group differences in *f*_P_ values will yield biased inferences when analysing *BP*_P_. Finally, *BP*_ND_ is one of the most common outcome measures in PET imaging, equal to 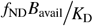, where *f*_ND_ represents the radioligand free fraction in the non-displaceable compartment. However, for this reason, group differences in *f*_ND_, will also yield biased inferences when analysing *BP*_ND_. These binding potential outcomes can be calculated directly from the rate constants of the PK model, which is known as direct estimation, however the reliability of these estimates has been found to be poor in many cases, particularly for *BP*_ND_ (Slifstein and Laruelle, 2001). Alternatively, binding potential outcomes can be calculated relative to a reference region in the brain which is devoid of specific binding, when one is available, under the assumption that non-displaceable binding is equal throughout the brain. This is known as indirect estimation, and can greatly improve the reliability of binding potential estimates (Slifstein and Laruelle, 2001). Indirect estimation of *BP*_ND_ is especially common as it does not require collection of arterial plasma samples, which is highly desirable as this procedure is costly, can be uncomfortable for research participants, and adds to the complexity of PET acquisition (Gunn et al., 2001). However, the improvements in reliability using indirect estimation come at a cost, since group differences in reference tissue binding will bias inferences. In summary, all binding potential outcomes are measures of specific binding relative to a reference concentration and should therefore be approximately proportional to one another in theory; however this is more complicated in practice as estimation inaccuracies or individual differences in *f*_P_, *f*_ND_ or reference-region binding can introduce bias for inference.

### 10.2 Supplementary Materials S2: Priors

#### 10.2.1 Priors for the SiMBA 2TC models

For all priors, we use the same terminology used in Matheson and Ogden (2022).

##### Global Intercepts

Note that all priors are defined over the natural logarithms of the parameters, where the mean is shown first and the standard deviation second.

For the SiMBA *BP*_F_ model, we used the following priors.

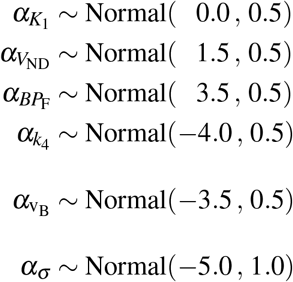

For the SiMBA *BP*_P_ and *BP*_ND_ models, we used the following priors.

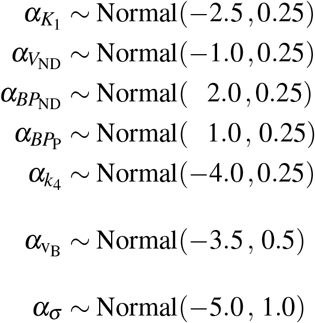

Hence forth all priors for *BP*_F_, *BP*_P_ and *BP*_ND_ are all the same. Hence I will write simply *BP*_X_.

##### Individual deviations

Differences between individuals were defined by specifying the primary pharmacokinetic parameters in one variance-covariance matrix, and v_B_ and *σ* on their own.

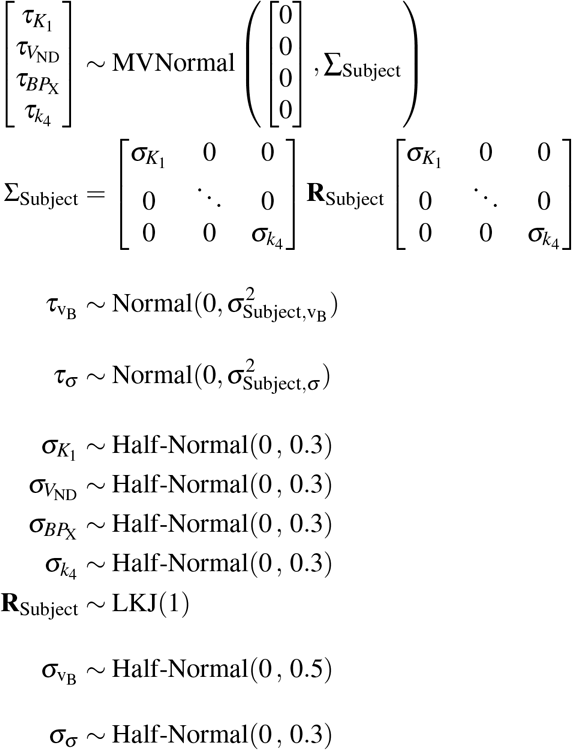

##### Regional deviations

For log *BP*_X_ and log *K*_1_, regional differences were defined as unpooled effects using a dummy (indicator) variable defined with reference to the dorsolateral prefrontal cortex as covariates. For simplicity, all regional differences were defined for all other regions as zero-centred regularising priors with the same SD.

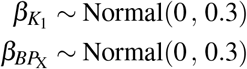

For the remaining parameters, regional differences were defined as partially-pooled variables, arising from a common distribution.

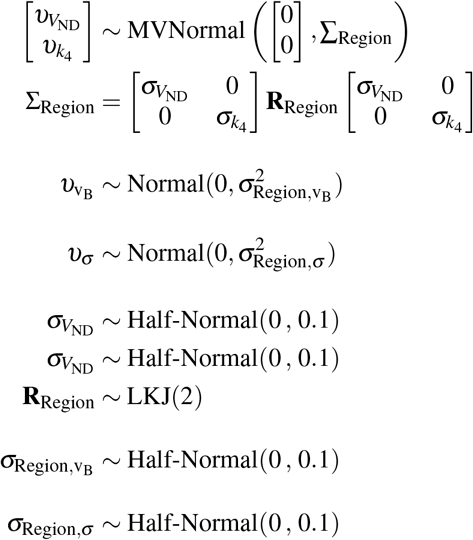

##### TAC deviations

Residual TAC effects were defined for the major four pharmacokinetic parameters, but were not included for v_B_ or *σ* as these were not considered to be of central importance.

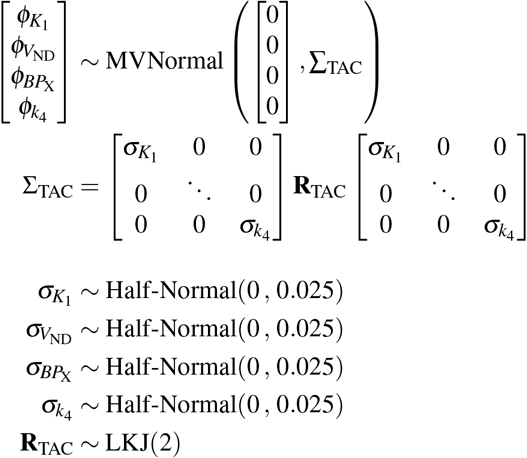

##### Covariates for PK Parameters

For the effects of group on BP, we defined priors for each region and diagnosis separately in order to detect regionally-specific effects.

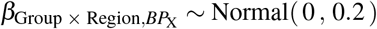

We also defined priors for age (per decade, centred) and sex as nuisance variables.

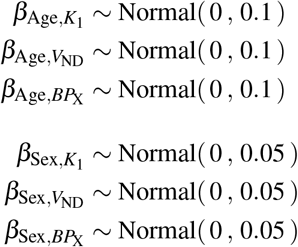

##### Measurement Error

For measurement error, *σ*, we used a combination of a global mean and partially pooled deviations across individuals and regions described above. We also made use of covariates for mean region size, injected radioactivity, and the duration of each frame, as well as a smooth function across the course of the PET measurement to accommodate any residual variability not accounted for by the covariates. The latter smooth term is expected to be more associated with the kinetics of the specific tracer after accounting for the influence of experimental factors.

All covariates are log-transformed and then centred. In this way, the global intercept is still interpretable as the geometric mean of the measurement error, and the covariates are related proportionally to the natural logarithm of the expectation of the measurement error.

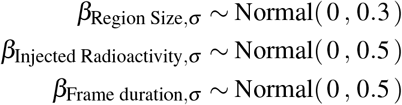

The smooth basis function makes use of a penalised regression spline. The *σ*_spline-coefficients_ term refers to the standard deviation of the spline coefficients, which penalises the wiggliness of the spline. The *α*_spline-coefficients_ term refers to the fixed effects term, which describes the magnitude of the influence of the smooth term around the estimated mean value. For the half-Student-*t* distributions, the first parameter represents the degrees of freedom, *ν*, followed by the location, *µ* and scale, *σ*, parameters.

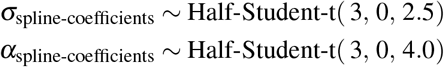

#### 10.2.2 Priors for the PuMBA SRTM model

For all priors, we use the same terminology used in Matheson and Ogden (2023).

##### Global Intercepts

Note that all priors are defined over the natural logarithms of the parameters, where the mean is shown first and the standard deviation second.

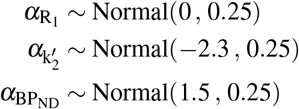

##### Individual deviations

Differences between individuals were defined by specifying the primary pharmacokinetic parameters in one variance-covariance matrix.

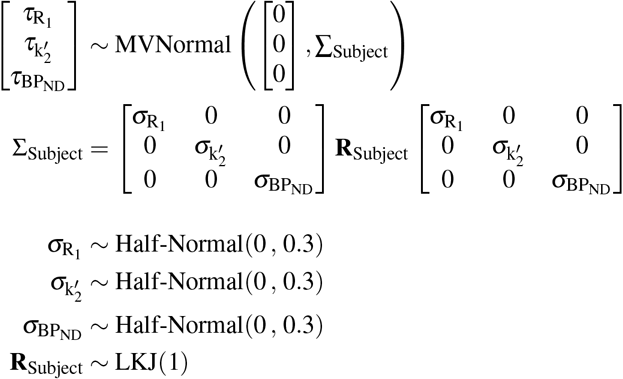

##### Regional deviations

For log *BP*_ND_ and log *R*_1_, regional differences were defined as unpooled effects using a dummy (indicator) variable defined with reference to the dorsolateral prefrontal cortex as covariates. For simplicity, all regional differences were defined for all other regions as zero-centred regularising priors with the same SD.

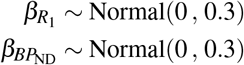

For 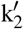, regional differences were defined as a partially-pooled variable, arising from a common distribution.

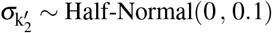

##### Covariates for PK Parameters

For the effects of group on *BP*_ND_, we defined priors for each region and diagnosis separately in order to detect regionally-specific effects.

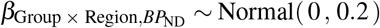

We also defined priors for age (per decade, centred) and sex as nuisance variables.

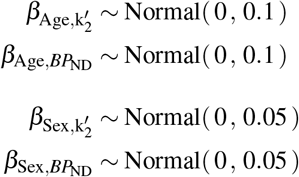

##### Residual deviations

We defined residuals, *ε*, also as being correlated across the PK parameters in the same way as the individual and regional deviations. The residual variance hence corresponds to a combination of the TAC deviations described above for the SiMBA models, as well as inaccuracy in the estimation of these parameters.

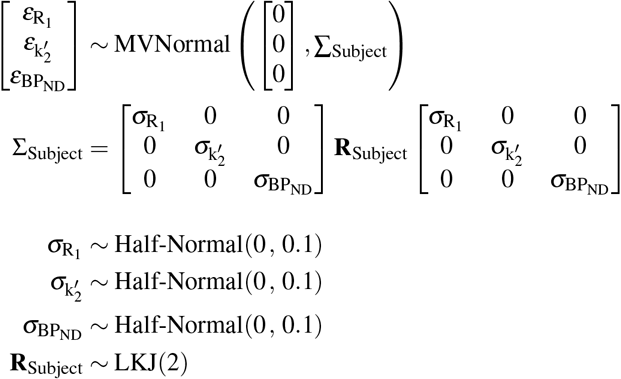

### 10.3 Supplementary Materials S3: Summary of Group Differences

#### 10.3.1 Fixed Regional Effects Results

Below are shown the regional differences as shown in Figure 2.

**Table S1.**
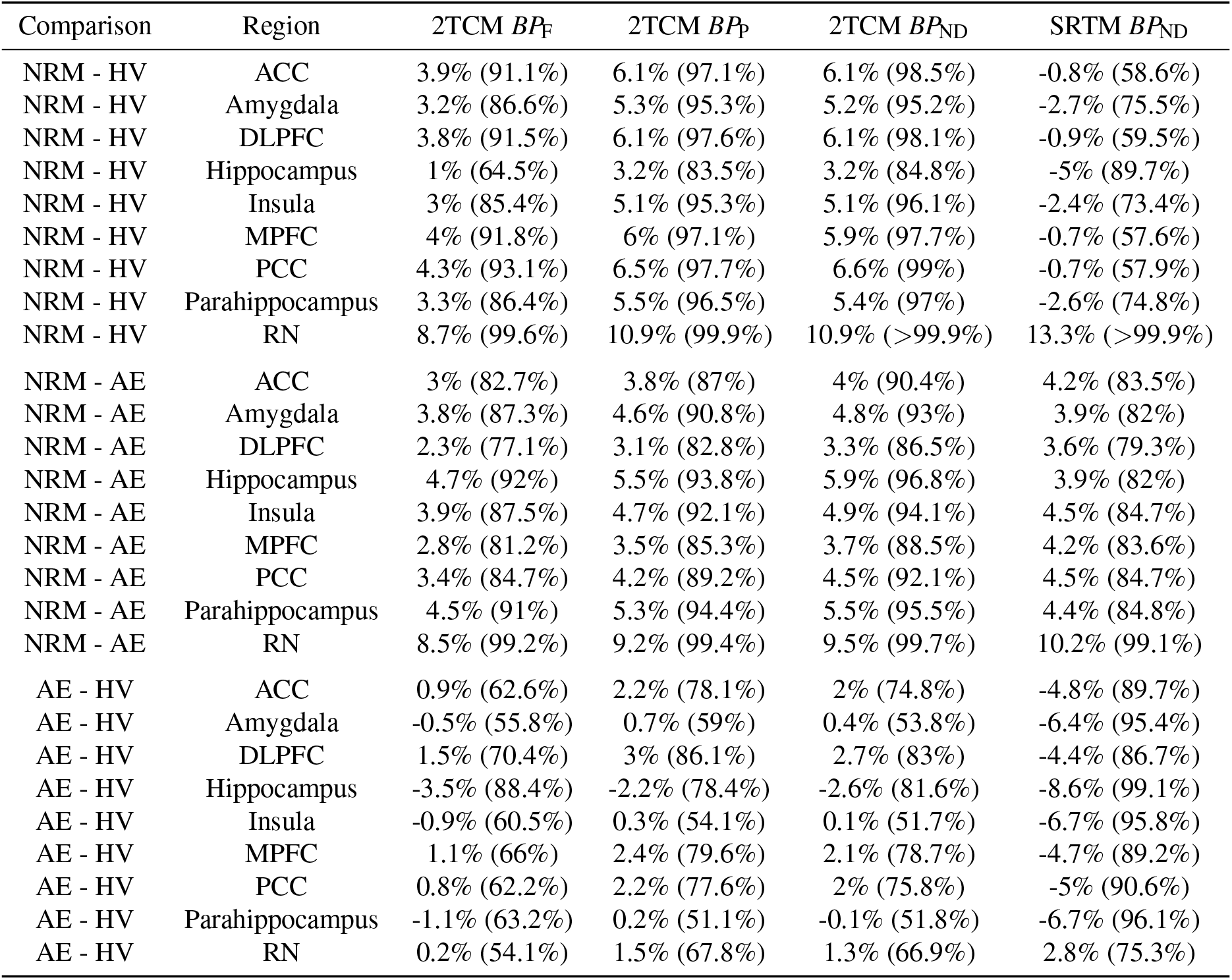
Summary of Group Difference Results: Estimate 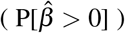

#### 10.3.2 Random Regional Effects Results

Below are shown the regional differences from the additional models fit with random slopes for regional differences in projection regions.

**Table S2.** Summary of Group Difference Results: Estimate 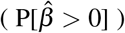

| Comparison | Region | 2TCM $BP_F$ | 2TCM $BP_P$ | 2TCM $BP_{ND}$ | SRTM $BP_{ND}$ |
| --- | --- | --- | --- | --- | --- |
| NRM - Control | Raphe | 9.6% (99.7%) | 11.1% (>99.9%) | 11.5% (>99.9%) | 11.6% (99.4%) |
| NRM - AE | Raphe | 8.7% (98.7%) | 9.8% (99.5%) | 9.9% (99.1%) | 10.2% (98.3%) |
| AE - Control | Raphe | 0.8% (59.1%) | 1.2% (63.6%) | 1.5% (66.2%) | 1.3% (60.9%) |
| NRM - Control | Projection | 4.2% (91.4%) | 5.8% (97.7%) | 6.2% (98.5%) | -3.4% (18.7%) |
| NRM - AE | Projection | 3.9% (90.2%) | 4.9% (95%) | 5% (95.5%) | 4.2% (85.7%) |
| AE - Control | Projection | 0.4% (54.5%) | 0.9% (62.1%) | 1.1% (65%) | -7.3% (2.6%) |

And below are the comparisons between the raphe and projection region differences between groups, i.e. comparing whether the group differences were statistically differentiably larger in the raphe nuclei compared to the average group differences in projection regions. Here the differences are presented in the units of logBP.

**Table S3.** Summary of Regional Differences in Group Difference Estimates

| Outcome | Group Comparison | Region Comparison | Difference ( $P[\hat{\beta} > 0]$ ) |
| --- | --- | --- | --- |
| 2TCM $BP_F$ | NRM - Control | Raphe - Projection | 0.050 (99.5%) |
| 2TCM $BP_P$ | NRM - Control | Raphe - Projection | 0.049 (98.9%) |
| 2TCM $BP_{ND}$ | NRM - Control | Raphe - Projection | 0.049 (99.3%) |
| SRTM $BP_{ND}$ | NRM - Control | Raphe - Projection | 0.050 (99.5%) |
| 2TCM $BP_F$ | AE - Control | Raphe - Projection | 0.004 (57.2%) |
| 2TCM $BP_P$ | AE - Control | Raphe - Projection | 0.003 (55.8%) |
| 2TCM $BP_{ND}$ | AE - Control | Raphe - Projection | 0.003 (54.4%) |
| SRTM $BP_{ND}$ | AE - Control | Raphe - Projection | 0.004 (57.2%) |

### 10.4 Supplementary Materials S4: Binding Potential Values

**Figure S1.**
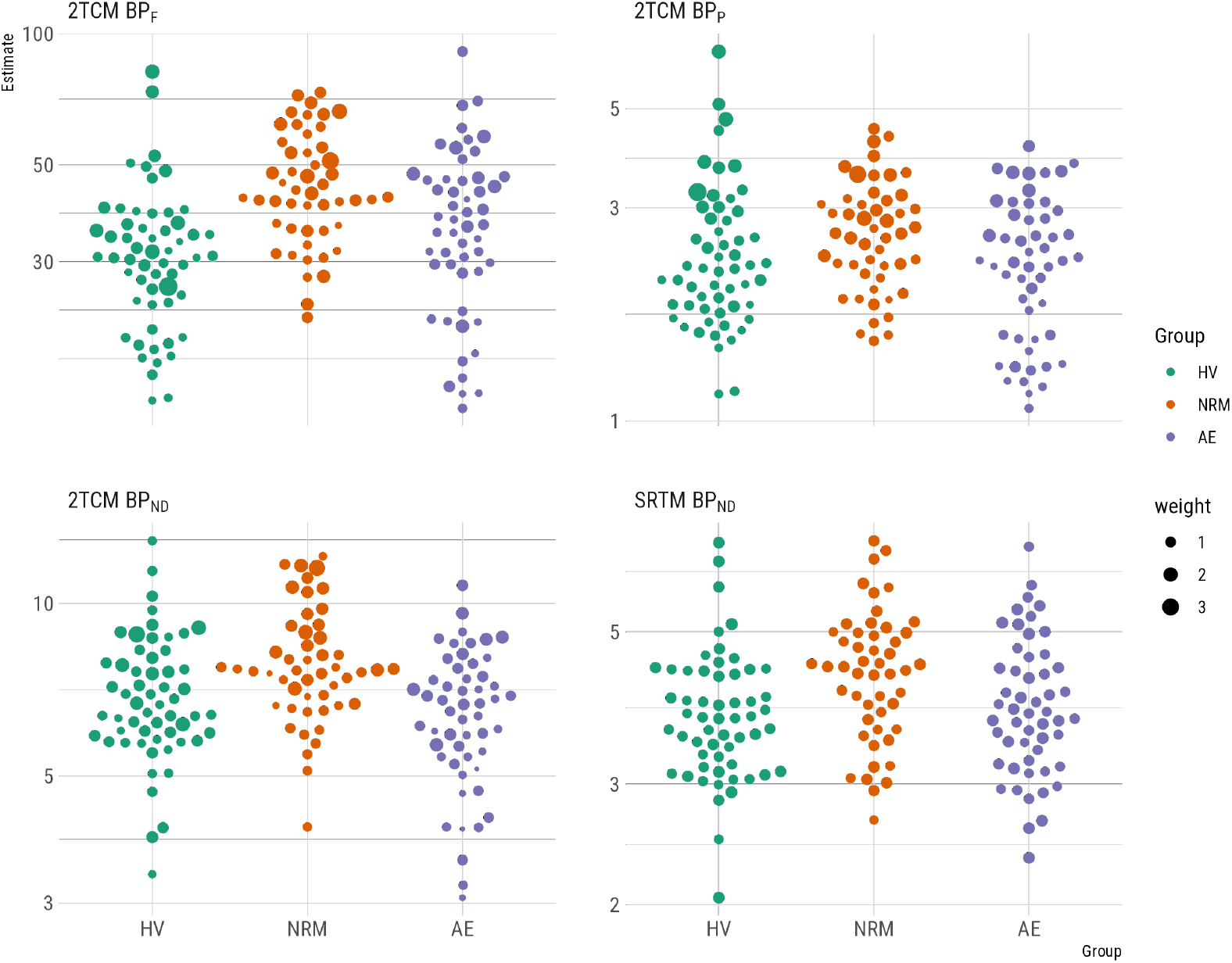
Raphe nuclei binding potential values for all outcome measures

